# Portello: Making global assembly more effective for rare-disease whole genome sequencing

**DOI:** 10.64898/2026.02.18.706714

**Authors:** Christopher T. Saunders, Zev Kronenberg, James M. Holt, William J. Rowell, Michael A. Eberle

## Abstract

**Motivation:** Long-read *de novo* assembly methods provide great potential to improve whole genome sequencing analysis for rare-disease, however these methods are typically under-utilized. Among the complications of an assembly-based approach is the difficulty of reviewing read evidence for assembly-based inferences, including access to read-based information on basecall qualities, methylation, and mosaic variants not captured in the assembly consensus. An additional complication is resolving conflicts in variant representation between assembly-based inferences and those based on conventional reference-based read-mapping pipelines. Here we propose a new assembly-based mapping approach, portello, which addresses both of these issues by transferring read mappings from the sample’s *de novo* assembly contigs onto a standard reference sequence.

**Results:** Portello’s assembly-based mapping approach results in read-to-reference alignments that can be used by standard mapping-based variant callers without modification, resulting in improved accuracy due to higher-quality mapping. This approach also enables assembly and reference mapping-based inferences to be unified onto a consistent view of the sample, and enables direct review of read-level support for the assembly consensus in the context of a standard reference genome. To demonstrate portello’s impact on variant calling, we show that DeepVariant calls from portello alignments remove 47% of small variant basecall errors found in calls made from conventional read mapping. We additionally demonstrate how portello alignments can improve visualization and interpretation of complex loci, including an example where a copy number gain in a segmental duplication region can be easily inferred from assembly-based mapping while interpretation of the same region from conventional read mapping remains impractical. Portello is also capable of directly phasing and haplotagging read alignments as part of the alignment transfer process, including for partially-phased assembly contig inputs where phasing blocks need to be delineated within each contig.

**Availability:** The portello source and pre-compiled binary is released on GitHub: https://github.com/PacificBiosciences/portello.

## Introduction

*De novo* genome assembly of HiFi long reads is a powerful approach to human rare-disease analysis, especially given that rare variants may be highly diverged from the reference genome^1^. Recent studies have demonstrated that a substantial fraction of candidate causal *de novo* and recessive variants can be found only with assembly-based variant detection^2^. In recent years, there have also been substantial advances in the speed and accuracy of assembly for diploid organisms^3–5^ which produce partially-phased/dual assembly outputs, or a haplotype-resolved diploid assembly when supplementary scaffolding information is available.

In spite of this improved sensitivity to detect causal variants, *de novo* assembly remains an under-utilized option for whole genome sequencing (WGS) analysis of rare disease. Several complications with assembly-based sequence analysis contribute to this phenomenon. When working with assembly contigs and assembly-based variant calls, it remains difficult to directly review the supporting read evidence of each variant call to assess artifacts and potential assembly errors. Furthermore, it is difficult to directly relate read-based information on basecall quality and methylation to the assembly-based variant calls, as well as detect and review mosaic variants that are unlikely to be found in the assembly contigs. An additional challenge is to synchronize the results of assembly-based variant calling with those from a conventional mapping-based variant calling pipeline. For users relying primarily on a mapping-based pipeline, it can be difficult to resolve potentially conflicting variant calls and representations of the sample between the two approaches. For instance, structural variant calling from assembly contigs might suggest a genomic region is deleted in the sample, but this region could still include many small-variant calls and smaller deletions from a mapping-based pipeline.

Here we describe a new assembly-based mapping approach that addresses these complications. This approach starts by generating a global *de novo* assembly, mapping reads back to the assembly contigs and then mapping the assembly contigs to any standard reference genome, such as GRCh38. Portello itself then transfers the read-to-contig alignments to the reference genome. This process creates assembly-based read mappings that can be used by any standard variant caller, yet retain consistency with assembly contig mappings. The portello approach improves read mapping through two complementary mechanisms. First, it incorporates long-range haplotype information captured in the assembly contigs, improving global read placement. Second, it increases local alignment accuracy by partitioning the alignment problem into read-to-contig alignment, dominated by sequencing error, and contig-to-reference alignment, dominated by sample variation. This separation yields more consistent read alignments on the reference genome. These improvements directly enhance variant-calling accuracy, enable clearer visualization and interpretation of read evidence, and support improved characterization of copy number variation.

While these variant calling improvements are a substantial benefit of assembly-based read mapping, the increased practicality of using an assembly-based workflow for rare-disease analysis is likely the most important improvement. With assembly-based read mapping, results from standard variant calling tools and assembly-based tools can be more easily merged into a consistent interpretation of the sample. Assembly-based read mappings also allow for read-level review and visualization of assembly-based variants, which can be viewed together with read-level data, for instance to understand a variant’s proximal methylation patterns.

## Results

### Improved read representation across the reference genome

Portello remaps reads from alignments on the sample’s assembly contigs to a standard reference genome, given the read-to-contig alignments and the contig-to-reference alignments as input (**Figure 1A**). This process effectively allows read alignment to be partitioned into the two simpler sub-problems of read-to-contig alignment where query differences are dominated by sequencing error, and contig-to-reference alignment where query differences are dominated by sample variation. The resulting remapped reads demonstrate more consistent local alignments in regions where sequencing error and biological variation can be difficult to partition, such as variable number tandem repeats (VNTRs), as well as improved mapping properties where the sample’s contigs can effectively act as sample-specific decoy sequence to remove extraneous mappings from reference regions. To demonstrate portello’s alignment output, read alignment views from an example VNTR region are shown for conventional mapping with pbmm2 (**Figure 1B**), compared to assembly-based mapping with portello (**Figure 1C**), demonstrating the higher variant representation consistency resulting from portello’s approach.

**Figure 1:**
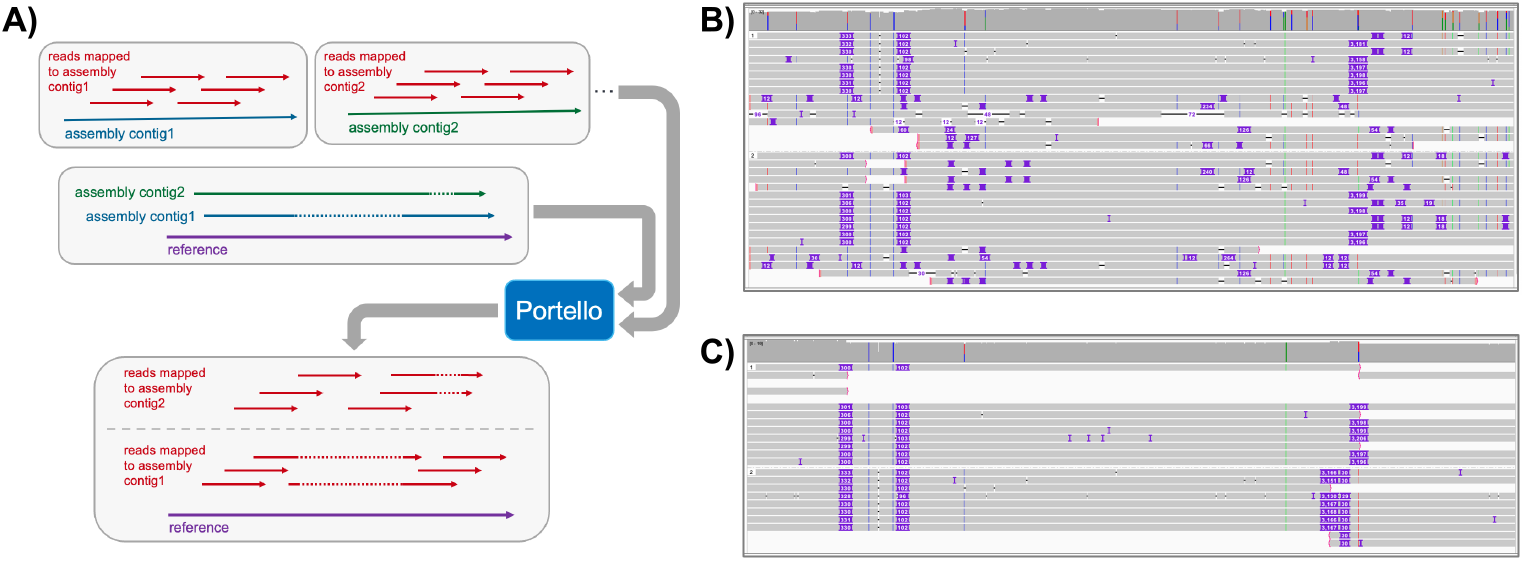
(A) Schematic demonstrating portello’s workflow, resulting in an assembly-based alignment of reads to a standard reference genome. A comparison of read alignments for an example VNTR region (HG002 chr4:40,294,825-40,295,700) shows (B) conventional mapping with pbmm2 vs (C) assembly-based mapping with portello. The same read data is used as input to both mapping processes so all differences are due to mapping methods.

### Small variant calling accuracy

Read remapping by portello results in a read alignment file that can be processed by variant calling tools designed for conventional long-read mapping to a reference genome. To systematically test this quality, as well as to better understand the genome-wide impact of the improved read representation example discussed above, we compare small variant calling accuracy from reads mapped by a conventional long-read mapper (pbmm2) vs. assembly-based mapping from portello. Using DeepVariant as an exemplar state-of-the-art long-read small variant caller, we call and assess small variants in two samples with high-quality small-variant benchmark sets from each of these read mapping processes, where the same set of reads is used in both the conventional (pbmm2) and portello mapping steps (see Methods). Our results from this evaluation show substantial variant calling accuracy improvement when DeepVariant starts from portello-remapped reads (**Figure 2, Table S1**). For both HG002 and NA12878 the primary impact is a reduction in false negatives, where portello removes 52% and 55% of pbmm2 false negative basecall errors in the two samples, respectively. This improvement in recall is accompanied by almost no change in false positive count. When considering total basecall errors (false positives and false negatives), portello removes 47% and 29% of all small variant basecall errors in the two samples, respectively.

**Figure 2:**
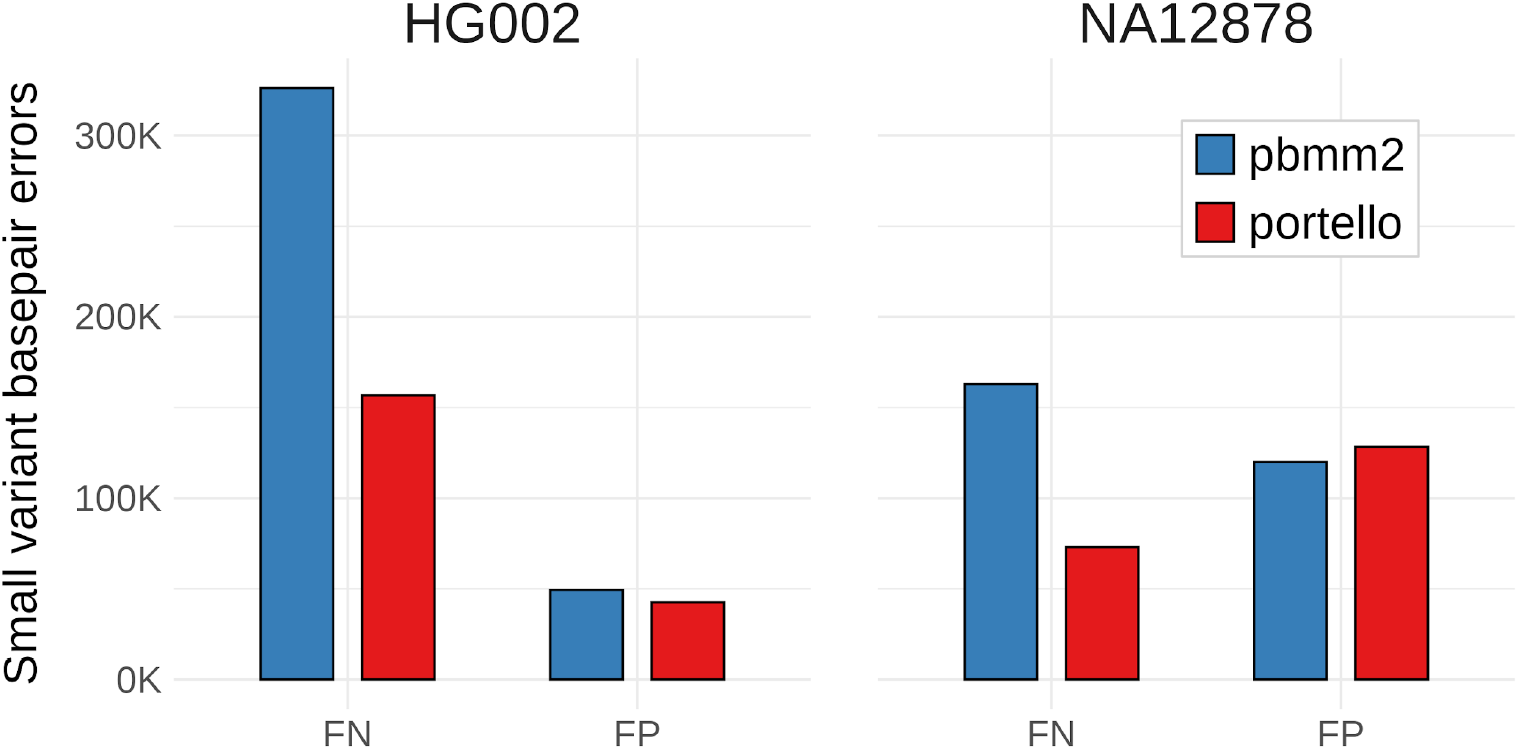
Comparison of DeepVariant small variant calling accuracy given either a standard read mapping input (pbmm2), or portello reads re-mapped from de novo assembly contigs. Comparisons are shown for two samples with high quality small variant benchmark sets, HG002 and NA12878. In both cases the same set of reads are available to the two mapping processes, and DeepVariant is run using the standard HiFi model without retraining for portello.

### Phasing accuracy

In addition to read remapping, portello also provides a novel method to annotate phasing on the remapped reads by leveraging the read-to-contig mapping. Portello provides two approaches to this annotation, depending on the type of input assembly contigs. When assembly contigs are haplotype-resolved (via parental sequencing, optical mapping, etc.), portello can simply assign phase to each read based on which contig it maps to in all reference regions with two distinct assembled contigs. Another common scenario is for the input assembly contigs to be partially-phased (i.e., dual assembly), in which case the contigs are expected to include haplotype switch errors. For this scenario, portello provides a simple variation on read-backed phasing based on the read-to-contig alignments, where the reads are evaluated for support across pairs of contig alleles that correspond to adjacent heterozygous variant locations in the contig-to-reference alignment.

We evaluate the accuracy of portello’s read phasing annotations for partially-phased assembly contigs by leveraging the draft GIAB T2T small variant benchmark for HG002. Direct evaluation of read haplotag accuracy vs. a small variant phasing benchmark is complicated, so an option was added to portello to output a phased VCF of heterozygous variants for benchmarking purposes. Whatshap^6^ is used to assess the VCF output over all autosomes and compare it with the T2T benchmark (see Methods). This evaluation shows that portello’s read phasing has a block NG50 of 334,357 bases, with a relatively low error count of 274 switches with 67 flips. With further development, there is significant potential to improve this capability in coordination with a portello integrated small-variant caller designed to more accurately call small variants from the read-to-contig alignments.

### Potential for improved CNV accuracy

To further demonstrate the diagnostic relevance of portello read remappings, we examine a copy number gain example drawn from the microarray-confirmed clinical CNV study by Gross et al.^7^ From this study we ran hifiasm^4^ assembly together with the full portello read mapping protocol on HiFi WGS data for NA21886, and created a standard mapping of the same reads using pbmm2 for comparison (see Methods). One of the array-based copy gains detected in this sample is a 225 kb duplication segment found in a segmental duplication region on chromosome 22. We examine this segment (lifted from hg19 to GRCh38) for the same set of NA21886 HiFi reads mapped by portello and by pbmm2. In this instance there are extreme coverage irregularities in the conventional pbmm2 mapping (**Figure 3A**), including a spike of nearly 12,000-fold coverage, followed by a coverage dropout over the genes in this region (DGCR6 and PRODH). By comparison, when viewing the portello mapping (**Figure 3B**), there is a 3-fold coverage region closely corresponding to the array segment, and the portello reads can be further grouped by assembly contig to simultaneously observe contig coverage over the region and assess whether the contigs represent a compression of the sample sequence, outlining a high-confidence CNV calling process enabled by portello read mapping. This case demonstrates the potential for substantially improved CNV calling by using assembly-based read mapping with portello.

**Figure 3:**
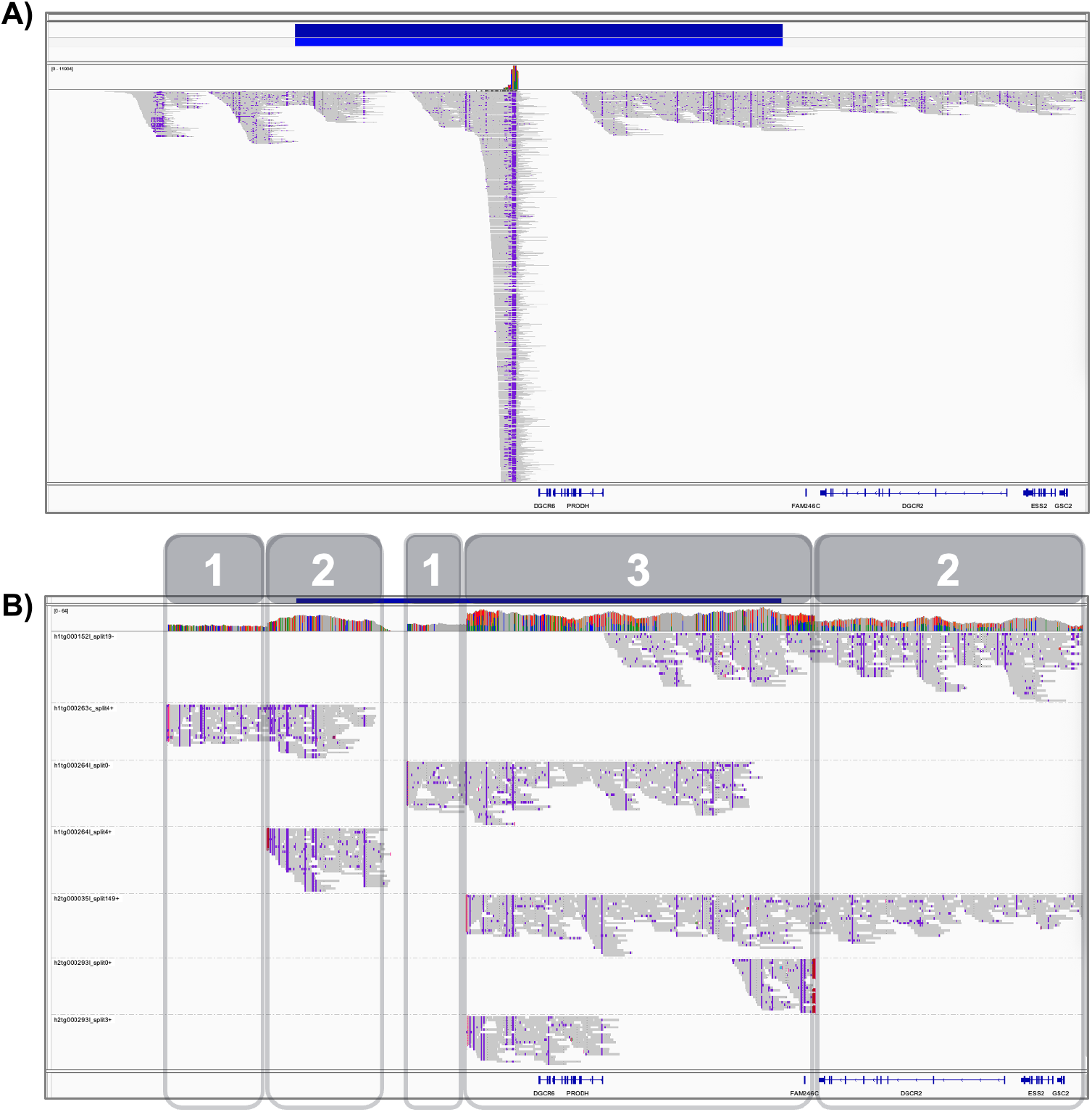
Read pileup views of a copy number gain in NA21886 confirmed by microarray in a previous clinical CNV study (Gross 2019). Both views show the IGV read pileup for the same region (chr22:18,680,093-19,158,500), but comparing mappings from pbmm2 (A) and portello (B), given the same read data input to each mapping process. The top bar in blue shows the microarray copy 3 region from Gross et al. In part B, the reads are grouped by the assembly contig segment each read maps to, using portello’s “ZS” read tag. Part B is additionally labeled with copy number segments, which can be inferred from portello alignments using both assembly contig depth and read coverage over each contig, here showing a copy 3 segment inference in a very similar region to the original microarray call.

## Methods

### Portello Methods

#### Read mapping transfer

To transfer read mappings from their own global *de novo* assembly contigs to a standard reference genome, portello relies on two primary alignment inputs. The first is alignments of reads back to their own assembly contigs. The second is alignments of assembly contigs to the standard reference genome. Given these inputs, portello creates a new alignment for each read by composition of the read-to-contig and contig-to-reference mapping functions, correctly accounting for any split alignment segments in both the read and contigs. Note although split alignment segments between the reads and their own assembly contigs are expected to be infrequent, these are important to correctly transfer read mappings for circular contigs such as the human mitochondria.

#### Pre-processing contig alignments

To improve the quality of the transferred read mappings, portello implements several pre-processing steps on the contig-to-reference mappings prior to read mapping transfer. These steps include (1) repeated matches trimming and (2) joining co-linear contig alignment mappings.

Repeated matches trimming is performed for each contig split alignment boundary such that each assembly contig base is mapped to the reference genome no more than once. For each instance of a repeated contig segment alignment to the reference, the alignment with the higher higher gap-compressed identity to the reference is retained and any other alignments are clipped. The recommended read-to-contig mapping procedure for portello uses a read mapper (pbmm2) which also implements repeated matching trimming. This combination means that when the recommended read mapping procedure is used, the transferred portello read mappings will retain the property that each read base is mapped to the reference genome no more than once – improving the ability to reason about genome coverage and copy number variation in downstream applications.

Joining collinear alignment segments from the contig to reference mapping is another pre-processing step which improves small variant calling accuracy of the remapped reads and makes the read alignments more comprehensible in downstream visualizations. When long assembly contigs are mapped to the reference genome by minimap2 (or similar mapping methods) the contig alignments are typically divided into multiple alignment segments by minimap2. In some cases the segmenting occurs even between two co-linear alignment segments wherever the read mapper’s Z-drop criteria are met, indicating a section of the contig that has greater local divergence from the reference sequence. Wherever these co-linear contig alignment segments are found, and the two contig alignments have the same mapping quality, portello will join them back into a single alignment segment. When joining the segments, any unmapped contig sequence intervening the two contig segments is annotated as an insertion rather than attempting to force an alignment. This segment joining procedure results in greater phasing contiguity and fewer “split” read alignments in the downstream read remapping process.

#### Left-shifting read alignment indels

Portello assumes that for both the read and contig alignment inputs, all alignments will follow the convention of left-shifting alignment indels. To retain this property in the transferred read mapping output, all read alignment indels must be left-shifted wherever any contig alignment segment is mapped to the reference in reverse orientation. This normalization is especially important for small variant call quality in regions of the reference genome where the two haplotypes are represented by contig segments mapped in opposite orientations, which would result in a mixture of left and right-shifted reads if this normalization were not applied.

#### Phasing annotation

In addition to transferring the read alignments to the reference chromosomes, portello also annotates phasing information for each transferred read mapping. The phasing annotation process follows the read ‘haplotagging’ conventions of methods such as whatshap^6^ and HiPhase^8^, wherein phase-annotated reads are given a haplotype identifier (1 or 2) using the bam auxiliary ‘HP’ tag, and a phase set identifier to annotate common phasing blocks using the bam auxiliary ‘PS’ tag.

The phasing annotation process begins while processing the assembly contig alignment segments. After reading in contig alignment segments and completing the repeated matches trimming and collinear segment joining steps, as described above, each contig alignment segment is annotated as a chromosome interval. These intervals are processed in sorted order by size from largest to smallest, where each interval is given a haplotype index based on how many previously processed intervals it overlaps. In portello, the criteria for the intervals to overlap is conservative. The intervals are considered to overlap if the intersection of the two intervals is greater than the minimum of 24 kb or one third of the new interval length. These haplotype identifiers are only annotated on reads where the total contig depth is two, and the read also has a phase set (see below).

The annotation process for phase sets depends on whether the input assembly contigs are being treated as fully-phased or partially-phased (as indicated from user input settings on the command-line). Assembly contigs should be treated as fully-phased when they’ve been scaffolded by longer-range phasing data from parental sequencing, optical mapping, etc. and can be approximated as representing a single sample haplotype. Partially-phased (or dual assembly) contigs from long-reads alone are assumed to contain switch errors.

In fully-phased contig mode, the phase sets are initially defined as the regions over which the same two contig alignment segments span a segment of the chromosome. The contig depth must be exactly two over these regions. The phase sets are refined to remove small interruptions in contig coverage, but merging adjacent depth 2 regions so long as the merged span is not more than 150% of the sum of component spans.

In partially-phased contig mode, the phase sets are determined by read support across heterozygous contigs as follows. First, in regions where the contig segment alignment depth is exactly two, heterozygous variants are approximated from the contig to reference alignments wherever at least one contig sequence varies from the reference and from the other contig. Next the read-to-contig alignments are evaluated to enumerate read support across each heterozygous variant location (in contig coordinates). Finally, phase-set blocks are extended through each adjacent heterozygous variant pair, as long as at least one read is found to support both variants.

In either mode, a read alignment segment is annotated with a phase set tag when its alignment intersects exactly one phase set region.

### Sample data

#### HG002

We used publicly available HiFi WGS data for HG002, sequenced to 32-fold coverage on the Revio system. The unmapped sequencing data can be downloaded from:

https://downloads.pacbcloud.com/public/revio/2022Q4/HG002-rep1/m84011_220902_175841_s1.hifi_reads.bam

#### NA12878

A cell line from NA12878 was sequenced to 34-fold coverage on the Revio system. The reads are available on the platinum pedigree S3 bucket at:

s3://platinum-pedigree-data/data/hifi/mapped/GRCh38/NA12878-cell-line-revio.GRCh38.haplota gged.bam

#### NA21886

This sample was ordered from Coriell and sequenced on the PacBio Sequel IIe platform with a target of 30-fold coverage.

### Conventional mapping

For all samples, the reads were mapped to GRCh38 with pbmm2 version 1.17.0 using the default “HIFI” preset.

### Portello assembly and mapping protocol

#### Assembly

For HG002 and NA21886, the sample input reads were provided as input to the HiFi-human-assembly-WDL pipeline version 1.0.2 installed from:

https://github.com/PacificBiosciences/HiFi-human-assembly-WDL

This pipeline uses hifiasm^4^ version 0.20.0 for contig assembly. The assembly was run to produce partially-phased (or dual-assembly) contigs which were used as input to the portello mapping protocol for each sample.

For NA12878 we used the Verkko diploid assembly from the recent Platinum Pedigree study^9^ available on S3 (s3://platinum-pedigree-data/assemblies). Although portello is primarily designed for the use case where the input reads are used for both *de novo* assembly and then for alignment back to their own assembly, in this case we demonstrate that the read remapping protocol is also valid when the sample assembly is produced from another source.

#### Assembly to reference mapping

For all samples, assembly contigs from both haplotypes were concatenated together and mapped to GRCh38 using minimap^10^ v2.30. The alignment command-line template used is:

~~~
minimap2 -L --secondary=no -a --eqx -x asm5 \
   -R “@RG\tID:{sample_id}\tSM:{sample}” \
   {reference_fasta} {assembly_contigs} |\
samtools sort - --write-index -o {sample}.asm.GRCh38.bam
~~~

Note that the ‘--eqx’ option to provide CIGAR strings with = and X match values is required for portello to work downstream. Additional information provided by the ‘--cs’ and ‘-Y’ options is not needed.

#### Read to assembly mapping

For all samples, assembly contigs from both haplotypes were concatenated together and used as the reference sequence for read alignment using pbmm2 v1.17.0 with the default “HIFI” preset.

#### Portello mapping

Given the contig to reference and read to contig mapping results, portello is then run for each sample as follows:

~~~
portello --ref {reference_fasta} \
    --assembly-to-ref {asm_to_ref_bam} \
    --read-to-assembly {read_to_asm_bam} \
    --remapped-read-output - \
    --unassembled-read-output unasm.bam \
    --phased-het-vcf-prefix remapped.{sample} \
    --input-assembly-mode partially-phased |\
samtools sort - --write-index -o remapped.{sample}.sort.bam
~~~

This configuration creates the primary portello bam output on GRCh38, together with a special phasing benchmark VCF that only includes phase-eligible heterozygous variants estimated from the contig to reference alignments. This file is used downstream to estimate the quality of the read phase set haplotags.

### Small variant accuracy assessment

For HG002 and NA12878, small variants are called from both the pbmm2 and portello read mapping outputs. Small variant calling is conducted with DeepVariant^11^ v1.9 using the standard HiFi model, i.e. the DeepVariant model was not trained with portello inputs.

Given the DeepVariant VCF output, small variant call quality was assessed using aardvark^12^ v0.10.3 according to the following template:

~~~
aardvark compare --reference {reference_fasta} \
  --truth-vcf {benchmark_vcf} --regions {benchmark_bed} \
  --compare-label {sample} --output-dir {sample} \
  --query-vcf {sample_deepvariant_vcf}
~~~

For HG002, aardvark was used with the GIAB draft small variant benchmark (V0.019-20241113) based on the T2T-HG002-Q100v1.1 diploid assembly aligned to GRCh38. The benchmark variants and confidence regions were downloaded from:

https://ftp-trace.ncbi.nlm.nih.gov/ReferenceSamples/giab/data/AshkenazimTrio/analysis/NIST_HG002_DraftBenchmark_defrabbV0.019-20241113/

For NA12878, aardvark was used with the platinum pedigree v1.2 truthset (s3://platinum-pedigree-data/truthset_v1.2/NA12878_hq_v1.2.smallvar.vcf.gz)

For both samples, aardvark’s ‘BASEPAIR’ comparison metric is used for all metrics and error counts discussed in this study.

#### Phasing accuracy assessment

Phasing accuracy was assessed for HG002 by using the same GIAB T2T small variant benchmark set used for small variant calling accuracy assessment.

As described above in the section on portello mapping, portello is capable of writing out a VCF of phase-eligible heterozygous variants, where these variants are detected directly from the contig to reference alignments. This VCF output is only intended to help benchmark the accuracy of portello’s haplotagging applied to the remapped read output, because accuracy assessment directly from the read haplotags can be substantially more difficult.

Using portello’s phased VCF output and the GIAB T2T benchmark we assess phasing quality using whatshap^6^ as follows:

~~~
whatshap compare --ignore-sample-name --tsv-pairwise compare.tsv \
  {benchmark_vcf} {portello_vcf}
whatshap stats --chromosome {autosome_list} --tsv stats.tsv {portello_vcf}
~~~

The total switch error count and phase block NG50 can then be taken from the compare and stats results, respectively.

## Conclusion

Assembly-based read mapping with portello has substantial potential to improve long-read rare-disease analysis. As demonstrated in this study there are immediate variant-calling accuracy gains, such as for small variants. We also show strong potential for new classes of hybrid variant callers integrating the contig and read inputs to portello, such as for CNV calling

where we show an example in which hybrid calling makes a simple and easily interpretable call which would be very difficult from conventional read mapping.

Perhaps the more important benefit of assembly-based read mapping is that it unifies the results of assembly-based and read-based inferences. With portello, one could use a contig-based SV caller, such as PAV^13^, to realize the higher accuracy of assembly-based structural variant calling, and combine this result with read-based tools for small variant calling on the portello remapped reads, so that all calls could be unified on a consistent view of the genome, for which the read evidence underlying all variant calls, including those made from assembly contigs, could be reviewed and visualized. In addition to these benefits, users will also acquire the ability to conveniently analyze mosaic variants, baseball qualities and methylation evidence in the context of their global *de novo* assembly results.

Given these benefits for rare-disease and other analysis scenarios, we anticipate the portello assembly-based read mapping approach should help users more easily and routinely leverage recent improvements in long-read diploid genome assembly, adding additional value to both existing and future long-read sequencing datasets.

## Acknowledgements

The authors thank Mitchell R. Vollger for early discussion on assembly-based read mapping.

## Supplementary Material

### Supplementary Tables

**Table S1:** Small variant calling assessment. For two samples, DeepVariant calls from pbmm2 and portello mapped reads were assessed against each sample’s respective small variant benchmark (see methods). All assessments were made with aardvark, and all analysis referenced in the manuscript use aardvark’s basepair comparison metric. Although the basepair metrics should generally provide the most accurate reflection of small variant calling performance, aardvark’s GT comparison metric is provided here for comparison.

| Comparison Metric | Sample | Mapping Process | FN | FP | Recall | Precision | F1 |
| --- | --- | --- | --- | --- | --- | --- | --- |
| BASEPAIR | HG002 | pbmm2 | 326253 | 49381 | 0.9830 | 0.9974 | 0.9901 |
| BASEPAIR | HG002 | portello | 156775 | 42593 | 0.9918 | 0.9978 | 0.9948 |
| BASEPAIR | NA12878 | pbmm2 | 163016 | 119992 | 0.9903 | 0.9929 | 0.9916 |
| BASEPAIR | NA12878 | portello | 73114 | 128362 | 0.9957 | 0.9924 | 0.9940 |
| GT | HG002 | pbmm2 | 47082 | 25366 | 0.9898 | 0.9945 | 0.9921 |
| GT | HG002 | portello | 32571 | 22118 | 0.9929 | 0.9952 | 0.9941 |
| GT | NA12878 | pbmm2 | 32500 | 33203 | 0.9925 | 0.9923 | 0.9924 |
| GT | NA12878 | portello | 21229 | 30008 | 0.9951 | 0.9931 | 0.9941 |

